# Proteomic Landscape of Human Spermatozoa: Optimized Extraction Method and Application

**DOI:** 10.1101/2022.11.25.518017

**Authors:** Mengqi Luo, Tao Su, Shisheng Wang, Jianhai Chen, Tianhai Lin, Qingyuan Cheng, Younan Chen, Meng Gong, Hao Yang, Fuping Li, Yong Zhang

## Abstract

Human spermatozoa proteomics exposed to some physical, biological or chemical stressors is being explored. However, there is a lack of optimized sample preparation methods to achieve in-depth protein coverage for sperm cells. Meanwhile, it is not clear whether antibiotics can regulate proteins to affect sperm quality. Here, we systematically compared a total of six different protein extraction methods based the combination of three commonly used lysis buffers and physical lysis strategies. The urea buffer combined with ultrasonication (UA_ultrasonication) produced the highest protein extraction rate, leading to the deepest coverage of human sperm proteome (5685 protein groups) from healthy human sperm samples. Since the antibiotics, amoxicillin and clarithromycin, have been widely used against *H. pylori* infection, we conduct a longitudinal study of sperm proteome via data-independent acquisition tandem mass spectrometry (DIA-MS/MS) on an infected patient during on and off therapy with these two drugs. The semen examination and morphological analysis were performed combined with proteomics analysis. Our results indicated that antibiotics may cause an increase in the sperm concentration and the rate of malformed sperm and disrupt proteome expression in sperm. This work provides an optimized extraction method to characterize the in-depth human sperm proteome and to extend its clinical applications.

## Introduction

Human spermatozoa (commonly called sperm), as very specialized cells with morphological and compositional differences distinct from those of other human cells, are produced by complex, intricate and tightly controlled spermatogenesis[1, 2]. Their major function is to deliver the paternal genomes, RNAs and even proteins to the egg[3]. Hence, defective or damaged sperm can disrupt sperm-egg binding or embryo development, even if the sperm genome is normal[4]. During spermatogenesis or ejaculated sperm storage, sperm are exposed to physical, biological or chemical stressors that could result in adverse changes in sperm motility, viability or acrosome status [5-7]. At the molecular level, changes in sperm quality are regulated by proteins[5]. Hence, it is of great significance to characterize the human sperm proteome exposed to different stressors to help us understand sperm biology and fertility issues.

As a new field in human reproduction studies, human sperm proteomics studies have revealed that sperm have complex protein compositions, shattering the old point of view that sperm have simple protein compositions[2]. These findings are mainly attributed to the rapid development of advanced mass spectrometry and proteomics research in recent years. Due to the unique structure and composition of sperm, great efforts have been made to reach a depth of proteome coverage in human sperm by improving the sample processing method, protein digestion strategy, peptide separation method, mass spectrometry and data analysis software[8]. In 2006, 2-DE and MALDI-TOF-MS analysis resulted in the identification of 98 proteins from human normozoospermic sperm[9]. The first large-scale analysis of the human sperm proteome (1,053 proteins) based on LC□MS/MS was reported in 2007[10]. A total of 4,675 human sperm proteins were identified using an LTQ Orbitrap Velos mass spectrometer in 2013[2]. In a more recent study in 2019, 4,959 human sperm proteins were identified (including 4,392 quantified proteins) using TMT-based LC□MS/MS[11]. In the same year, 5,246 human sperm proteins were identified, and 3,790 proteins were quantified using advanced Orbitrap Fusion mass spectrometer[12]. In addition, Castillo *et al*. reviewed relevant studies in sperm proteome on the basis of RNA-seq and protein database, and summarized 6,871 human sperm proteins, which play important roles in fertilization, early embryo development, and modulating gene expression [13]. These studies provide abundant information about human sperm protein compositions. However, compared to the conventional cell and tissue proteomic studies, there is a lack of systematic exploration of protein extraction methods for the special sperm cells. We reasoned that the optimized methods and combined with advanced mass spectrometer techniques would enable more high-confidence human sperm proteins to be identified and quantified.

Male infertility factors might be indicated using the traditional semen analysis screening[14]. However, LC□MS/MS-based proteomics information can provide vital clues toward elucidating human sperm protein function at the molecular level, including fertilization and development[15]. Understanding the impact of the sperm proteome on the developing embryo may help to promote eugenics and develop diagnostic and therapeutic methods for male infertility. Several previous studies have focused on the human sperm proteome from asthenozoospermic patients or failure of in vitro fertilization[16-19]. Meanwhile, a few studies have revealed the significant proteome alteration in sperm exposure to some external factors, such as cryopreservation, heat stress, extremely low electromagnetic field (ELEF), agonists, ejaculatory abstinence periods, and reactive oxygen species etc. which could affect the quality of the ejaculated sperm[6, 7, 11, 12, 18, 20, 21]. In addition, the posttranslational modifications (phosphorylation, glycosylation, etc.) of human sperm proteome are being explored[20, 22, 23]. Antibiotics are one of the most widely used clinical drugs that resist pathogenic microorganisms[24]. However, with the overuse of antibiotics, the growing antibiotic resistance of microorganisms has become a serious global problem affecting human health, such as reproductive disorders[25]. For example, inflammation in the epididymis and testis caused by bacteria contributes significantly to male infertility[26]. This has led to a growing demand for novel antibiotics and calls for an end to antibiotics misuse[27]. However, whether antibiotics can directly affect sperm protein composition and function is still unclear. Some *in vitro* studies suggest that antibiotics (amikacin, penicillin, gentamicin, etc.) can be safely added to semen extenders to control the growth of bacteria contaminating ejaculated mammalian semen without affecting sperm quality (motility, acrosomal, plasma membrane integrity, etc., but may contribute to the development of antibiotic resistance[28, 29]. However, the influences of antibiotics on human sperm proteome upon oral during infection have not been well studied.

*Helicobacter pylori* (*H. pylori*) is a gram-negative bacterium in the human stomach and has been classified as a group 1 carcinogen. *H. pylori* has an overall prevalence of 44.3% worldwide. China has the largest number of infected populations with with an infection rate of over 50%. This infection can cause gastric atrophy, peptic ulcer disease and gastric cancer[30]. The current recommended first-line *H. pylori* eradication antibiotics are amoxicillin and clarithromycin[31]. Hence, there is a large male population that takes both amoxicillin and clarithromycin. However, the impact of on the sperm quality after taking amoxicillin and clarithromycin remains elusive.

In this study, we systematically optimized the sperm protein extraction methods and constructed an in-depth human sperm proteome database using advanced proteomics techniques and mass spectrometry. Taking advantage of the optimized methods, we combined quantitative proteomics approaches together with semen examination and morphological analysis to conduct a longitudinal study in a patient who is taking amoxicillin and clarithromycin due to *H. pylori* infection. The sperm proteome was analyzed and compared in on and off medication conditions. Our results suggest that the abnormal regulation events occurring in human sperm proteins may play a significant role in sperm quality and reproductive potential after amoxicillin and clarithromycin therapy.

## Experimental Procedures

### Materials and chemicals

Phosphate-buffered saline (PBS), dithiothreitol (DTT), iodoacetamide (IAA), Tris base, acetone, and urea were purchased from Sigma (St. Louis, MO, USA). Formic acid (FA), trifluoroacetic acid (TFA), acetic acid (HAc), acetonitrile (ACN), ethanol (EtOH) and methanol (MeOH) were purchased from Merck (Darmstadt, Germany). C8 and C18 reverse-phase medium were purchased from Agela Technologies (Tianjin, China). The 30-kDa centrifugal filters were purchased from Merck Millipore (Carrigtwohill, Ireland). Sequencing grade trypsin was obtained from Promega (Madison, WI, USA). Sephacryl was purchased from Affymetrix (Santa Clara, CA). BluPower Fast Staining Coomassie was purchased from Zoonbio (Nanjing, China). The quantitative colorimetric protein and peptide assay kits were purchased from Thermo Fisher Scientific (Waltham, MA, USA). Deionized water was prepared by a Milli-Q system (Millipore, Bedford, MA). All other chemicals and reagents of the best available grade were purchased from Sigma□Aldrich or Thermo Fisher Scientific.

### Biospecimen collection

Human semen was collected from healthy donors in West China Second University Hospital of Sichuan University, Sichuan Province, China. This study was approved by the Ethics Committee of West China Second University Hospital of Sichuan University and abided by the Declaration of Helsinki principles. Conventional semen analysis was conducted in accordance with guidelines from the World Health Organization Laboratory Manual for the examination and processing of human semen. Semen examination and morphological analysis were performed by West China Second University Hospital of Sichuan University for proteomic analysis. The semen samples from ten healthy donors met the following parameters: pH≥7.2, sperm concentration≥15*10^6/mL, sperm survival rate≥58%, round cell<1*10^6/mL, forward motile sperm (A+B)≥32%, and normal sperm percentage≥4%. They were centrifuged at 4 °C (2000 g×20 min) to separate spermatozoa from seminal plasma. Spermatozoa were washed three times using PBS buffer (pH 7.4) to remove residual seminal plasma. Spermatozoa were collected in 1.5 mL tubes and stored at –80 °C until use. A young man (23 years old) with Helicobacter pylori infection was treated with antibiotic therapy (amoxicillin 2 g/day for 15 days, clarithromycin 1 g/day for 15 days). During the medication (2022/3/29∼2022/4/11), semen samples were taken approximately every 3 days for a total of 5 times. After drug withdrawal (2022/4/12∼2022/4/28), semen samples were also taken approximately every 3 days for a total of 5 times. These semen samples were produced by masturbation and collected into sterile sample containers. Semen samples were preprocessed and stored as above. We collected written informed consent from all volunteers.

### Protein extraction

The obtained spermatozoa samples from ten healthy controls were pooled and divided. An equal number of spermatozoa were placed in 2 mL tubes and resuspended using 500 μL of RIPA buffer (50 mM Tris-HCl (pH 7.4), 150 mM NaCl, 1% Triton X-100, 1% sodium deoxycholate, 0.1% SDS), UA buffer (100 mM Tris-HCl (pH 8.5), 8 M urea), or SDS buffer (100 mM Tris-HCl (pH 7.6), 2% SDS). Then, each sample was processed by ultrasound (Kunshan, China) and/or a homogenizer (Hangzhou, China). The ultrasound instrument is set as power = 100 W; frequency = 25 KHZ; total time = 15 min; on = 7 s; off = 3 s. The homogenizer is set as speed = 6.5 m/s; time = 3 min; number of cycles = 3. The processed samples were centrifuged at 13,000 × g for 15 min at 4 °C. The supernatant from samples treated in different solutions and ways was collected and stored at −80 °C until use. The total protein concentration was determined by using a Bradford or BCA protein assay kit.

### SDS□PAGE analysis

SDS□PAGE analysis was performed as follows: a constant volume of supernatant was boiled 10 min after adding 5× SDS□PAGE loading buffer. After centrifugation at 13,000 × g for 15 min, the samples were loaded onto 12% SDS□PAGE gels. Electrophoresis was begun at 100 V at 4 °C. After 20 min, the voltage was set at 120 V, and the time was set at 1 h. After the dye front had run off the gel, the lanes were cut out and washed with destaining solution (25% (v/v) ethyl alcohol, 8% (v/v) acetic acid, 67% (v/v) water) for 30 min. BluPower Fast Staining Coomassie was used for dyeing.

### Reduction, alkylation and digestion

Spermatozoa proteins (100 μg) were proteolyzed using a filter-aided sample preparation (FASP) protocol. Briefly, precooled acetone was added to the supernatant containing RIPA buffer or SDS buffer at a ratio of 3:1 (V/V) at −20 °C for 4 h. Then, the spermatozoa proteins were precipitated and collected after centrifugation at 2,000 × g for 10 min at 4 °C. The proteins were freeze dried and suspended in 100 μL of 8 M UA buffer. All samples in UA buffer were added to a 30-kDa filter. After centrifuging at 13,000 × g for 15 min at 25 °C, 200 μL of 8 M UA buffer with 20 mM DTT was added, and the reduction reaction was carried out for 4 h at 37 °C. Then, 50 mM iodoacetamide (IAA) was added and incubated in the dark for 1 h at room temperature. UA buffer was replaced by 50 mM ammonium bicarbonate buffer by centrifugation at 13,000 × g for 15 min. Two micrograms of trypsin was added to each filter tube, and the proteins were digested at 37 °C overnight. The filter tubes were washed twice with 100 μL of water by centrifugation at 13,000 × g for 15 min. The flow-through fractions were collected. The peptide concentration was measured using a quantitative colorimetric peptide assay. The peptide mixtures were freeze-dried and then stored at −80 °C.

### Prefractionation of peptides

The peptides were redissolved in 90 μL of ammonia water and separated using high-Ph reversed-phase chromatography. In brief, a pipet tip (AXYGEN, USA) containing a layer of C8 membrane and 5 mg of C18 reverse-phase medium was washed with 90 μL of methanol and ammonia water (pH 10.0), respectively. Fifty micrograms of peptides were loaded onto the tip and centrifuged at 1,200 × g for 5 min. The peptides were eluted with increasing concentrations of acetonitrile in ammonia water (6, 9, 12, 15, 18, 21, 25, 30, 35, and 50%, v/v). The 10 fractions were collected and defined as F1 (6% + 25%), F2 (9% + 30%), F3 (12% + 35%), F4 (15%), F5 (18%), and F6 (21% + 50%). These fractions were freeze-dried and stored at −80 °C.

### Western blot analysis

The protein concentration of each sample was determined by the BCA method (Pierce, Rockford, USA). After their electrophoretic separation on 12% SDS□PAGE gels, these protein samples were transferred onto polyvinylidene difluoride membranes and blocked with 5% nonfat dry milk (NFDM). Membranes with protein samples subsequently reacted with anti-ACTB (1:5000, pAb, ZEN BIO, China), anti-SCRN2 (1:500, pAb, ZEN BIO, China), anti-FDFT1 (1:500, mAb, ZEN BIO, China) and anti-PLPP1 (1:500, pAb, ZEN BIO, China) overnight at 4 °C and then HRP-conjugated secondary antibody (1:2000, ZEN BIO, China) for 1 hour at room temperature. Actin (ACTB) was used as an internal control. Both chemiluminescent visualizations were performed using an ECL detection system.

### LC-MS/MS analysis

All samples were analyzed by LC-MS/MS using an Orbitrap Fusion Lumos Mass Spectrometer (Thermo Fisher, USA). Specifically, peptides were dissolved in 0.1% FA and separated on a 100-μm-inner-diameter column with a length of 25 cm (ReproSil-Pur C18-AQ, 3 μm; Dr Maisch) over a 78-min gradient (buffer A, 0.1% FA in water; buffer B, 0.1% FA in 80% ACN) at a flow rate of 400 nl/min. In addition, 2 μL of iRT reagent was added to the samples for DIA analysis.

The parameters for DIA analysis were as follows: (1) MS1: Orbitrap resolution = 60,000; scan range (m/z) = 350–1500; RF lens = 40%; AGC target = custom; maximum injection time = 50 ms; (2) MS2: Orbitrap resolution = 30,000; HCD collision type = 30%; first mass (m/z) = 150; RF lens = 40%; AGC target = custom; maximum injection time = 54 ms; The parameters for DDA analysis were as follows: (1) MS: Orbitrap resolution = 60,000; scan range (m/z) = 350–1500; RF lens = 40%; AGC target = custom; maximum injection time = 25 ms; included charge state = 2–7; exclusion after n times, n = 1; exclusion duration = 18 s; (2) MS2: Orbitrap resolution = 15,000; isolation window (m/z) = 1.6; HCD collision type = 30%; first mass (m/z) = 150; AGC target = custom; maximum injection time = 22 ms.

### Data analysis

The DIA raw data files were searched against the human UniProt database (UP000005640_9606) using Spectronaut (version 15.4.210913.50606). The parameters were as follows: the analysis type was “directDIA”. The fixed modification was carbamidomethyl (C). Variable modifications included acetyl (protein N-term) and oxidation (M). All other settings were set at the default values.

The DDA raw data files were searched against the human UniProt database (version 2015_03, 20,410 entries) using MaxQuant (version 1.5.3.8). The search parameters were set as follows: Two missed cleavage sites were allowed for trypsin digestion. Carbamidomethyl (C) was set as a fixed modification. Oxidation (M) and acetyl (protein N-term) were set as variable modifications. All other settings were set at the default values.

### Bioinformatic analysis

Data preprocessing, statistical analysis, and data presentation were processed in our in-house platform (https://www.omicsolution.org/wkomics/main/). Data are reported as the mean ± standard deviation. Quantified proteins with missing value ratios above 30% across all samples were removed, and imputation was implemented with the k-nearest neighbor (KNN) algorithm based on the median normalized protein abundances. Student’s t test was used for statistical analysis. Proteins with p values less than 0.05 were considered significant. Gene Ontology (GO) analysis and Kyoto Encyclopedia of Genes and Genomes (KEGG) pathway analysis were performed on the DAVID website (https://david.ncifcrf.gov/).

## Results

### Comprehensive comparison of the identified human spermatozoa proteins using different sample processing methods

To compare and evaluate different human sperm protein extraction methods, we collected semen samples from 10 healthy donors and obtained pooled sperm by centrifugation. Equal amounts of sperm cells were resuspended using a constant volume of lysis buffer (UA, RIPA, or SDS buffer). After extraction using ultrasonication or homogenization, sperm proteins can be obtained by centrifugation (**Fig. 1A**). Hence, a total of six different protein extraction methods (UA_ultrasonication, UA_homogenization, RIPA_ultrasonication, RIPA_homogenization, SDS_ultrasonication, SDS_homogenization) were compared. SDS□PAGE analysis results show that most human sperm proteins can be extracted by these methods. However, there were significant differences in the type and abundance of extracted proteins (**Fig. 1B**). To determine which extract proteins are different, these sperm proteins were digested into peptides and analyzed by DIA mass spectrometry, which has the characteristics of high quantitative accuracy and high reproducibility, and Spectronaut software[32]. Three technical runs from each method were performed for subsequent analyses, and each run resulted in different protein identifications, ranging from 4309 to 4447. A total of 4470 protein groups could be quantified, 4302 (∼96.2%) protein groups were common to all six methods, and some protein groups were only identified in a single method (**Fig. 1C** and **supplemental Table S1**). Compared with the UA_homogenization, RIPA_ultrasonication, RIPA_homogenization, and SDS_homogenization methods, the protein and peptide identifications in the UA_ultrasonication (4,443±7 proteins and 37,911±41 peptides) and SDS_ultrasonication (4,438±6 proteins and 37,829±126 peptides) methods achieved deeper coverage (*p*<0.05, Student’s T test). However, there was no significant difference in protein and peptide identification numbers between the UA_ultrasonication and SDS_ultrasonication methods (*p*>0.05, Student’s T test) (**Fig. 1D** and **supplemental Table S1**). These results suggested that the type and number of proteins extracted by different methods are different.

**Figure 1.**
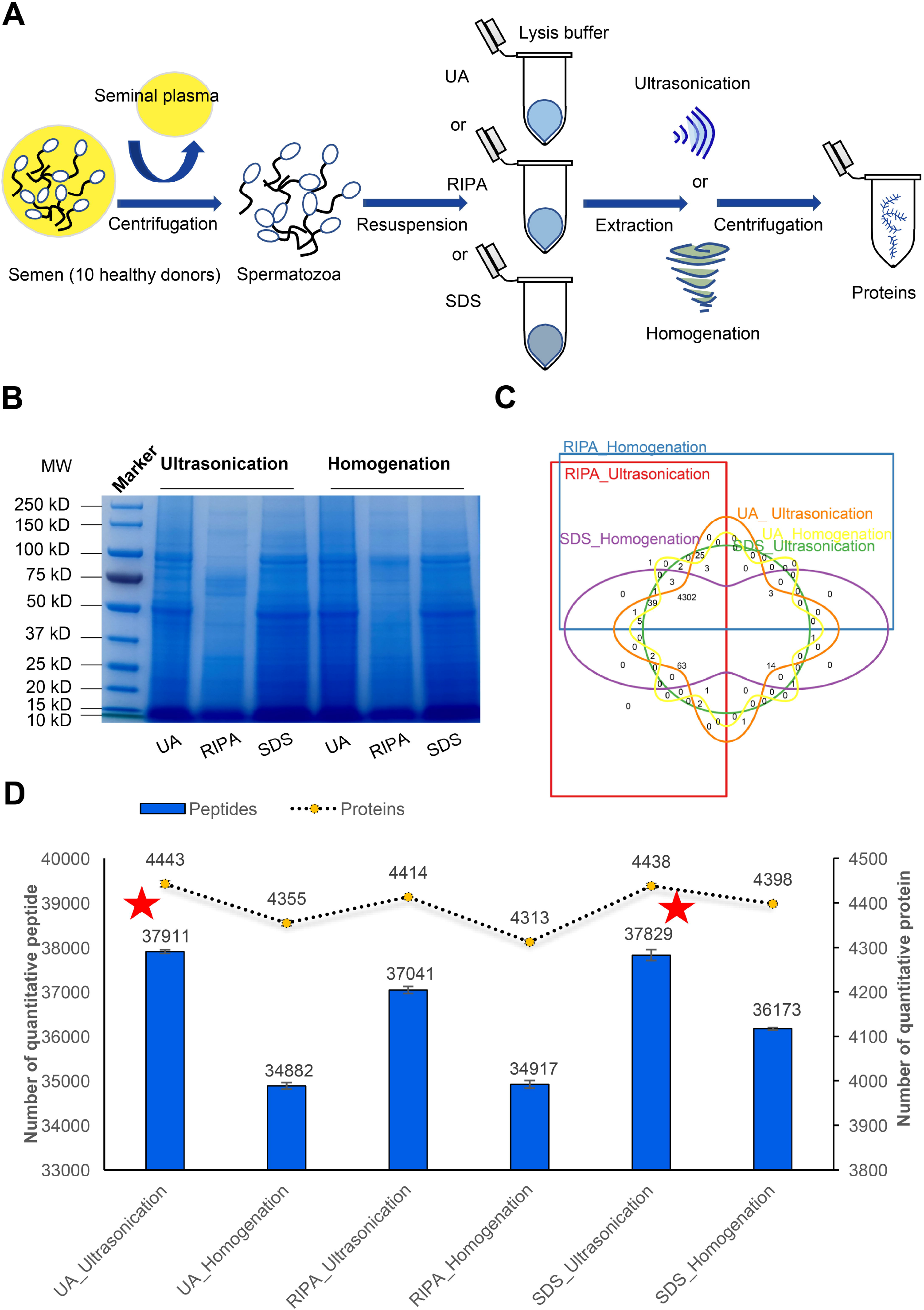
Comparison of quantitative proteins using different sample processing methods. **A**, Workflow of semen sample processing; **B**, SDS□PAGE analysis of sperm proteins; **C**, Venn plot of quantified proteins; **D**, Number of quantitative peptides and proteins.

To further analyze the protein quantified differences extracted by UA_ultrasonication and SDS_ultrasonication methods, quantitative protein difference analysis was carried out. As shown in **Supplemental Fig. S1 and Table S2**, 444 of 4323 protein groups (384 were increased and 60 were decreased in the UA_ultrasonication method) were significantly different (p<0.05, fold changes>2 or <0.5). This showed that using the UA_ultrasonication method can result in more kinds of protein to obtain a higher protein extraction rate. GO analysis showed that these proteins were mainly located in the extracellular exosome and cytoplasm cytosol and were involved in spermatogenesis and ATP binding (**supplemental Fig. S2**). In addition to the advantage mentioned above, we used the UA_ultrasonication method in subsequent experiments when considering that UA_ultrasonication is advantageous for both soluble and hydrophobic proteins and compatible with FASP and LC□MS/MS.

### In-depth human sperm protein profiling

For in-depth human sperm protein profiling, we carried out the following experimental design. As shown in **Fig. 2A**, human sperm proteins were extracted using the UA_ultrasonication method and digested using trypsin. The peptide products were separated using C18 high-pH reversed-phase chromatography. After combining into 6 fractions, they were analyzed via DDA mass spectrometry with higher quality fragmentation information and MaxQuant software. Finally, 5685 unique protein groups with 46,562 peptides and 83,580 matched MS2 spectra were identified (**Fig. 2B and supplemental Table. S3**). To the best of our knowledge, this is the largest human sperm database known to date. The sperm proteins identified in this study were compared with previously published sperm proteome data in Shen *et al*.*’*s study article (5,246 proteins) and Castillo *et al*.’s review article (6,871 proteins)[11, 13]. In the current study, 4868 identified proteins covered published sperm proteome data previously, and 817 other proteins were newly characterized in human sperm (**Fig. 2C**). GO analysis of these sperm proteins showed that they were mainly located in the cytosol and were involved in the protein transport process and protein binding function (**Fig. 2D**). Therefore, based on the optimized sperm protein extraction method and advanced mass spectrometry method, in-depth human sperm profiling can be realized. Knowledge of the human sperm proteome is essential for understanding reproductive functions.

**Figure 2.**
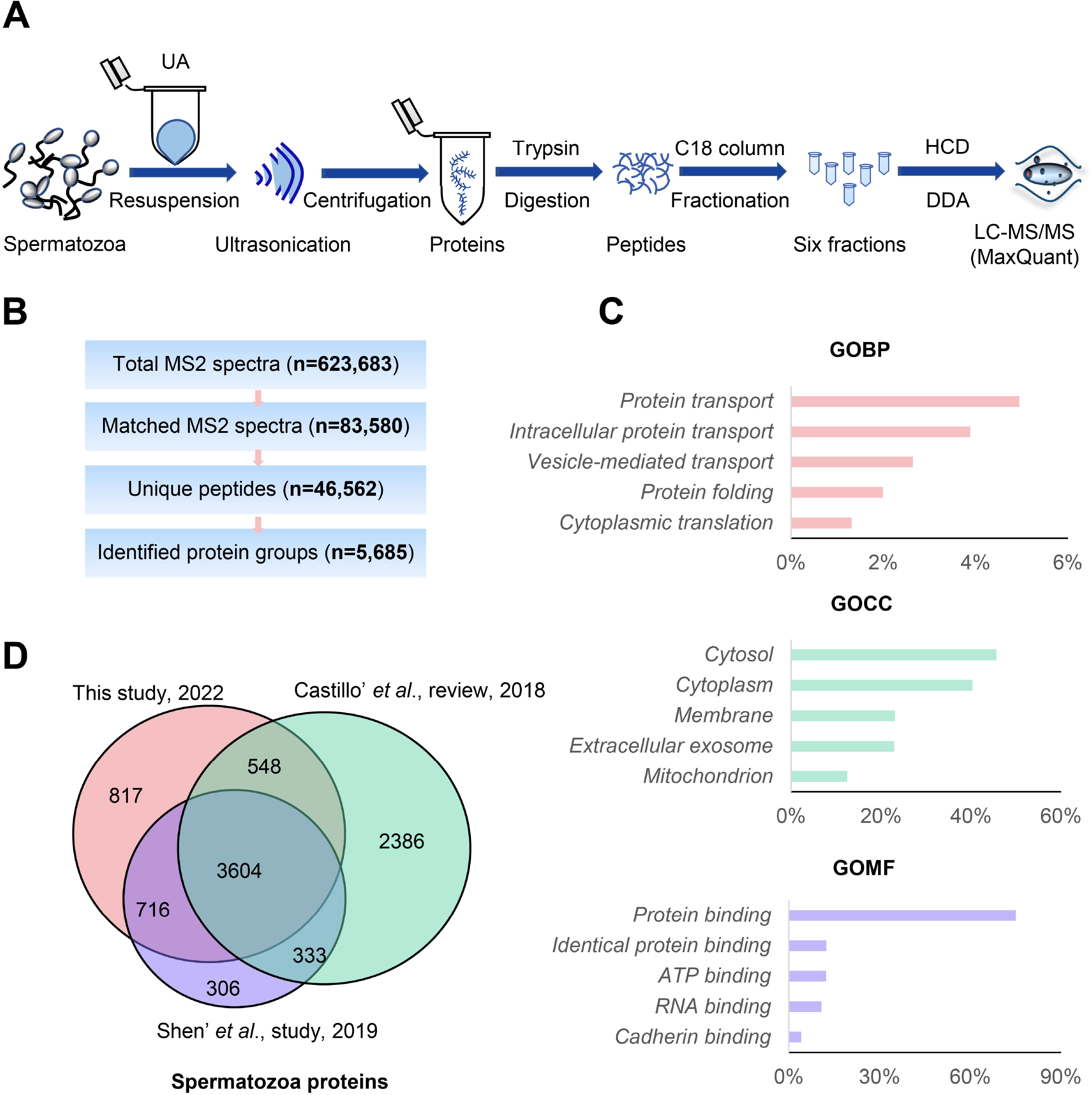
In-depth proteomics analysis of human sperm. **A**, In-depth proteomics workflow of human sperm based on DDA; **B**, Analysis summary of LC□MS/MS spectral database search: total MS2 spectra, matched MS2 spectra, unique peptides and identified protein groups; **C**, Gene ontology analysis of identified proteins; **D**, Sperm protein groups identified currently compared with the previous study and review.

### Changes in human spermatozoa proteins after antibiotic therapy

Semen examination and morphological analysis of human sperm at the medicine (a combination of amoxicillin and clarithromycin) period (5 sperm samples) and withdrawal period (5 sperm samples) were performed (**Fig. 3 and supplemental Table. S4**). The morphologically normal sperm has been progressively redefined as having a normal head (with acrosome), midpiece, and tail [33]. As shown in **Fig. 3A and supplemental Table. S4**, over 200 spermatozoa per sample were analyzed by light microscopy after being stained via a modified Papanicolaou method. The “normal” head of sperm should have an oval shape with smooth contours; its acrosome should be clearly visible and cover 30–60% of the anterior portion of the sperm head; its midpiece should be axially attached to the head, less than 1 μm in width and approximately 1.5 times the head length; its tail should be apically inserted to the postacrosomal end of the midpiece, approximately 45–50 μm long[34]. If the morphological analysis does not match the above descriptions, the sperm is defined as an abnormality or malformation. The sperm concentration of the ten samples collected in the medication period (M) and withdrawal period (W) are shown in **Fig. 3B**. The result shows that taking amoxicillin and clarithromycin may cause human sperm densities to rise (from 19.90± 1.32*10^6/ml to 75.89±4.76*10^6/ml). When the antibiotics were stopped, sperm densities returned to lower levels and leveled off. Furthermore, we compared the ratio of morphological abnormalities of the sperm head, midpiece, and tail between the M and W groups (**Fig. 3, C and D**). The results showed that the ratio of sperm with midpiece or tail defects increased significantly in the M group (p<0.05, Student’s T test) (**supplemental Table. S4**). This implies that taking amoxicillin and clarithromycin may cause an increase in the rate of malformed sperm in humans.

**Figure 3.**
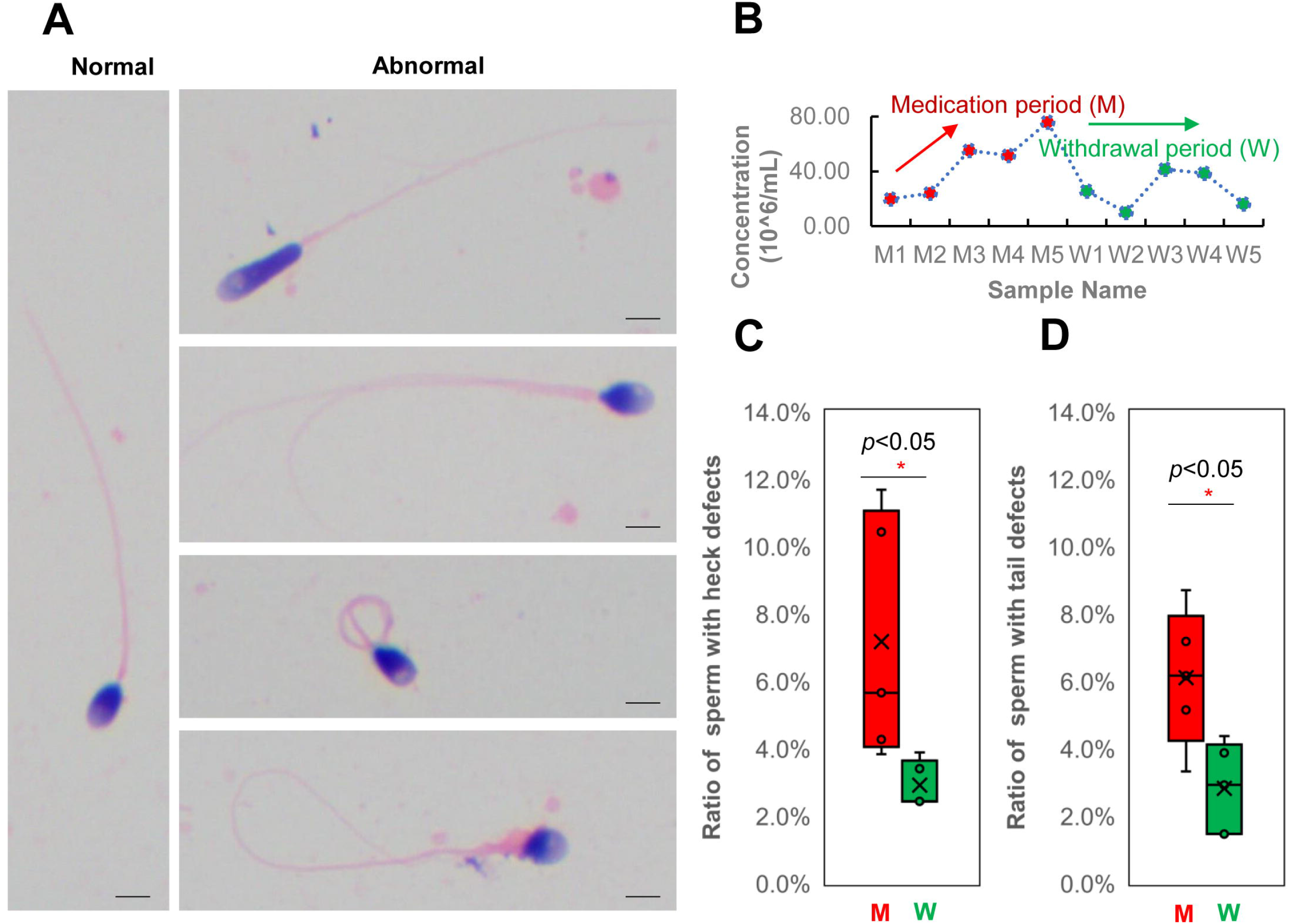
Semen examination and morphological analysis of human sperm after antibiotic therapy. **A**, Abnormalities in sperm morphology observed in sperm after antibiotic therapy by the Papanicolaou method and light microscopy (scale bars, 5 μm); **B**, Sperm concentration of the ten samples collected in the medication period (M) and withdrawal period (W); **C**, Ratio of sperm with heck defects in the M and W groups; **D**, Ratio of sperm with tail defects in the M and W groups.

To gain insight into the protein molecular changes in sperm, comparative proteomics analysis of the two groups was performed. As shown in **Fig. 4A**, the sperm samples from the M group (n=5) and W group (n=5) were processed using the optimized UA_ultrasonication method and analyzed in triplicate by the direct DIA approach and Spectronaut software. Then, statistical analysis was performed to identify differentially expressed proteins (**Fig. 4B and supplemental Table. S5**). A total of 5455 protein groups were quantified from the two groups. 5361 protein groups with less than 30% missing quantification values were retained. After normalization, missing values were filled by KNN algorithm and protein groups with CV less than 0.3 were retained. A total of 2413 protein groups were analyzed statistically. Student’s t test resulted in 255 differentially expressed proteins between group M and group W at p<0.05. For practical purposes, the candidates were further narrowed down to 159 protein groups (139 were upregulated and 20 were downregulated in group W) by ≥1.25-fold or ≤0.80-fold (**Fig. 4C**). Principal component analysis (PCA) of the 159 differentially expressed protein groups showed that these sperm proteins can separate the two groups (**Fig. 4D**). GO analysis showed that they were mainly located in the cytosol and were involved in the phosphorylation process and ATP binding function (**Fig. 4E**). The quantification information of these proteins is shown in **Fig. 5**. Two clusters were found, and most differentially expressed proteins (139/159) were downregulated in growing M. This implies that the combination of amoxicillin and clarithromycin can inhibit most sperm protein expression. The results were further confirmed by western blot. In **Fig. 6A**, two unchanged proteins (ACTB and SCRN2) and two upregulated proteins (FDFT1 and PLPP1) were quantified via western blot. The quantitative results were consistent with those obtained by mass spectrometry (**Fig. 6B**). In summary, combined amoxicillin and clarithromycin therapy may disrupt proteome expression and regulate metabolic pathways.

**Figure 4.**
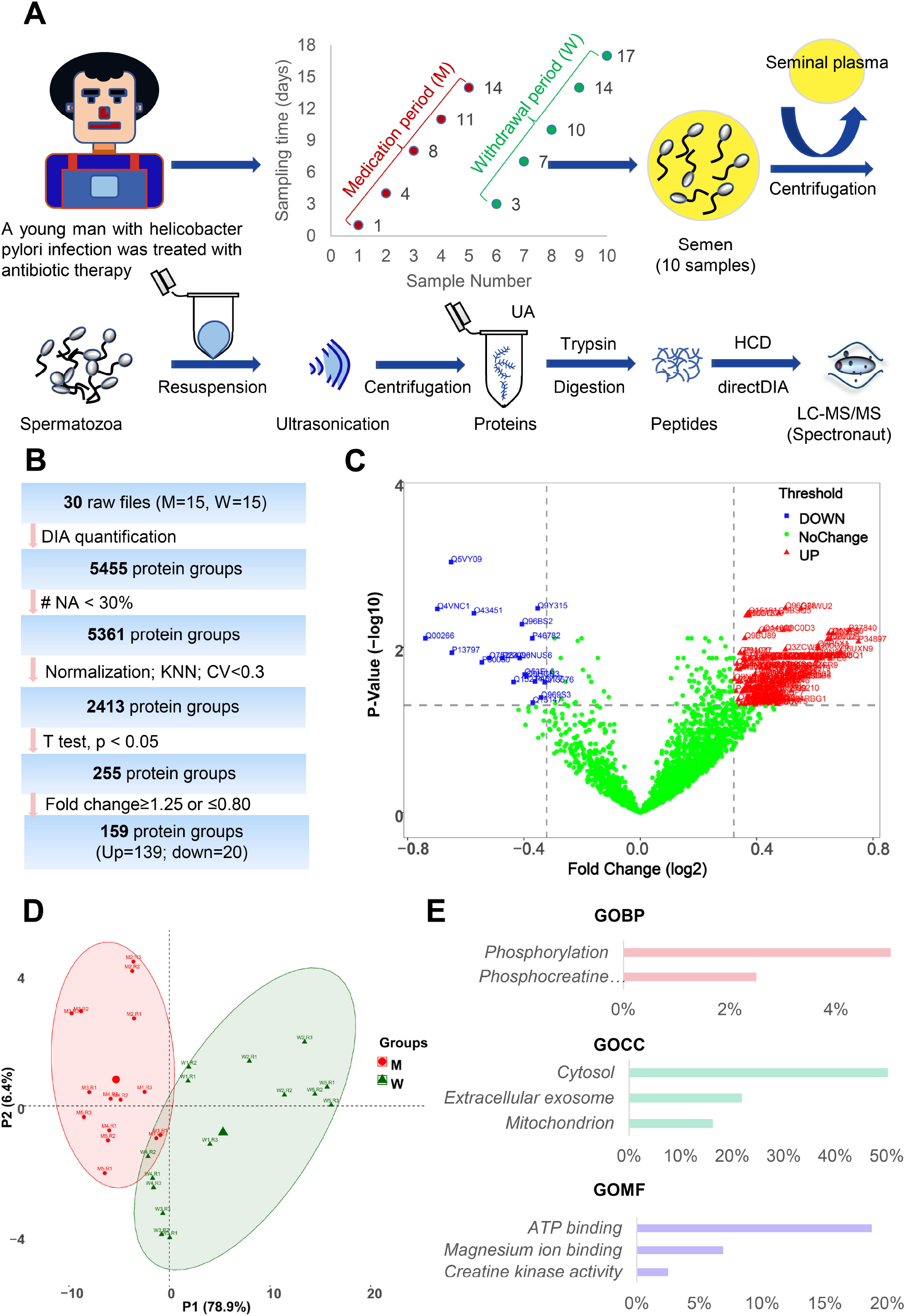
Quantitative proteomics analysis of human sperm after antibiotic therapy. **A**, Quantitative proteomics workflow of human sperm based on direct DIA; **B**, Analysis summary of LC□MS/MS spectral database search using strict thresholds; **C**, Volcano plot of sperm proteomic data. **D**, PCA of differentially expressed sperm proteins; **E**, Gene Ontology analysis of differentially expressed sperm proteins in the W group compared to the M group.

**Figure 5.**
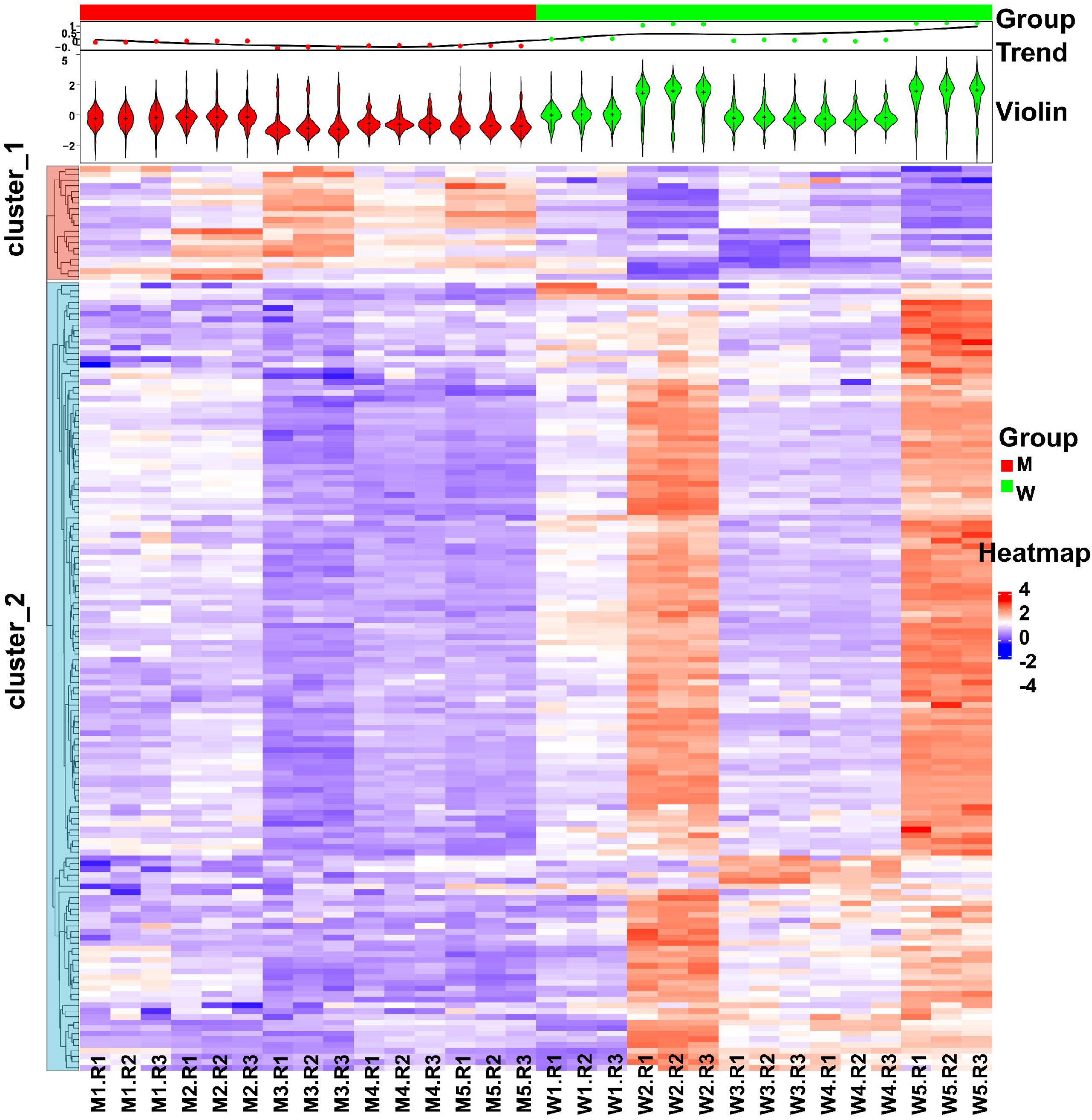
Heatmap analysis of differentially expressed sperm proteins between the M and W groups.

**Figure 6.**
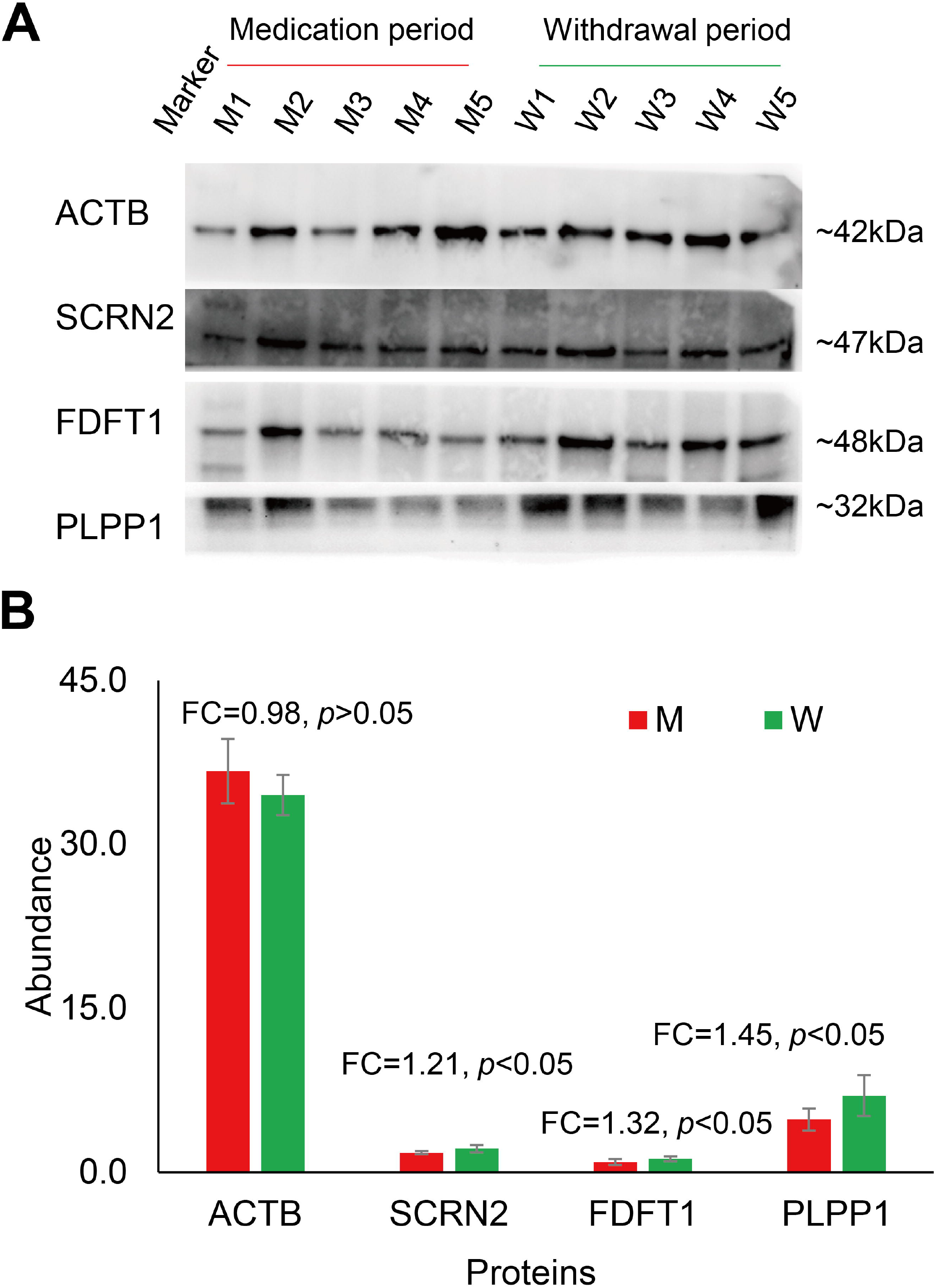
Western blot validation (A) and LC□MS/MS quantification (B) of four representative sperm proteins.

## Discussion

Protein extraction is a key analytical step in the study of proteomics because inadequate extraction will result in the loss of qualitative information and the deviation of quantitative results[35]. The commonly used buffers for protein extraction are SDS and UA buffer, which readily solubilize most proteins in different clinical samples[36, 37]. Commonly used extraction methods include ultrasonic instruments and homogenate instruments, which promote sample fragmentation and protein dissolution[38, 39]. The selection of an appropriate buffer and extraction instrument will help us extract adequate proteins for precise qualitative and quantitative analyses in clinical samples. However, a systematic comparison of human sperm proteome extraction methods is still lacking[40]. In this study, a total of six different protein extraction methods were compared. Both UA_ultrasonication and SDS_ultrasonication can achieve deeper protein and peptide identification (Figure 1). The reason may be that human sperm, as a kind of special cell, are better suited to using ultrasound instruments to extract proteins. Homogenate instruments are more suitable for extracting protein from enough tissue samples rather than microscale sperm cells because the homogenate beads will cause more protein residue. In addition, SDS must be removed completely from the sample prior to LC□MS/MS analysis because it can form salt crystals to clog the LC columns. Therefore, the UA_ultrasonication method was chosen as an optimized method for human sperm protein extraction in this study. Many buffers and extraction methods suitable for protein extraction have been reported. Each buffer or method was favorable for a certain subset of proteins. However, only the most commonly used methods were systematically compared in this study. Hence, in the future, researchers can explore more combinations of extraction buffers and means to increase the number of human sperm proteins identified.

With the development of MS-based proteomics technology and software, more diverse sample proteomes and deeper proteomes can be achieved[41, 42]. The recently reported size of the human sperm proteome was 5,246 identified proteins and 3,790 quantified proteins based on LC□MS/MS[12]. In addition, a total of 6,871 unique human sperm proteins have been reviewed[13]. It would be interesting to find new sperm proteins and achieve in-depth human sperm proteome coverage using an optimized sperm protein extraction method and advanced LC□MS/MS. As shown in Figure 2, 5685 unique protein groups with 46,562 peptides and 83,580 matched MS2 spectra can be identified. The size is the largest human sperm proteome. Such a good result is due to the optimized sperm protein extraction method, peptide fractionation, and advanced mass spectrometry. It is worth noting that we only used pooled sperm cells from healthy donors. Some specific proteins expressed in some patients or particular conditions could not be identified in this study. Large cohorts of samples from different sources will expand the scope of the sperm proteome.

In China, over 50% of people have been infected with gastric *Helicobacter pylori* (*H. pylori*), which causes gastric atrophy, peptic ulcer disease and gastric cancer[30]. Combined amoxicillin and clarithromycin therapy is the recommended first-line H. pylori eradication antibiotic[31]. This means that there is a large male population that takes both amoxicillin and clarithromycin in China. However, few studies have examined the effects of combined amoxicillin and clarithromycin therapy on human sperm in vivo. Hence, we performed a longitudinal study of the sperm proteome of a young man treated with combined amoxicillin and clarithromycin therapy. Five samples were collected during the medicine period, and five samples were collected during the withdrawal period. Semen examination and morphological analysis suggested that amoxicillin and clarithromycin therapy may cause an increase in the human sperm densities and the rate of malformed sperm (Figure 3). Comparative proteomic results further revealed 159 protein groups differentially expressed in the treatment period (Figure 4 and Figure 5). These results showed that amoxicillin and clarithromycin can disrupt proteome expression in human sperm and regulate metabolic pathways. In addition, to verify the accuracy of the quantitative results of mass spectrometry, we used a biochemical method (WB) to verify some specific proteins. Beta-actin (ACTB) is a highly conserved and high-abundance protein that polymerizes to produce filaments that form cross-linked networks in the cytoplasm of cells[43]. Antibodies against beta-actin are useful as loading controls for WB. Secernin-2 (SCRN2) is a protein located in extracellular exosomes. The function of this protein is not clear, and it is predicted to be involved in proteolysis[44]. Based on the LC□MS/MS quantification results, there was no significant difference in the expression of the two proteins between the M and W groups. The WB results also showed that the results were consistent (Figure 5). Farnesyl-diphosphate farnesyltransferase 1 (FDFT1) is a protein located in the endoplasmic reticulum membrane and the first committed enzyme of the sterol biosynthesis pathway. Phospholipid phosphatase 1 (PLPP1) is a protein that catalyzes the dephosphorylation of a variety of glycerolipid and sphingolipid phosphate esters and is involved in the regulation of inflammation and may regulate phospholipid-mediated signaling pathways[45]. Based on the LC□MS/MS quantification results, there was a significant difference in the expression of the two proteins between the M and W groups. The WB results also showed that the results were consistent (Figure 5).

The advantage of this work was the first comprehensive human sperm proteome study to explore the effect of amoxicillin and clarithromycin on sperm proteomics. However, it is difficult to continuously collect sperm samples from different patients. Hence, a limitation of this study was the limited number of patients. Additionally, the biological functions of differently expressed proteins in this study didn’t been studied. Given our results, we propose alteration of human sperm proteins may be associated with taking antibiotics. Antibiotics may be avoided when it is necessary to preserve sperm quality for reproductive purpose. Nonetheless, the suggestion needs more experimental and clinical evidence to verify.

## Conclusions

In this study, we proved that the urea buffer combined with ultrasonication (UA_ultrasonication) can produce the highest protein extraction rate, leading to the deepest coverage of human sperm proteome (5685 protein groups) from healthy human sperm samples. Taking advantage of the optimized methods, we combined quantitative proteomics approaches together with semen examination and morphological analysis to conduct a longitudinal study in a patient who is taking amoxicillin and clarithromycin due to *H. pylori* infection. The semen examination and morphological analysis indicated that antibiotics may cause an increase in the sperm densities and the rate of malformed sperm. The sperm proteome was analyzed and compared in on and off medication conditions. Our results suggested that the abnormal regulation events occurring in human sperm proteins may play a significant role in sperm quality and reproductive potential after amoxicillin and clarithromycin therapy.

## Acknowledgments

We would like to express our special thanks to the Human Sperm Bank of West China Second University Hospital for providing human semen samples. We thank the Wukong platform for providing data analysis support.

## Data Availability

The raw MS data have been deposited to the ProteomeXchange Consortium via the PRIDE partner repository with the dataset identifier **PXD036023**.

## Competing Interests

The authors declare that they have no competing interests.

## Author Contributions

Y.Z. and F.L. directed and designed the research; Q.C. performed semen examination and morphological analysis. Y.Z., M.L., T.S. and H.Y. directed and performed analyses of mass spectrometry data; Y.Z., J.C. and S.W. adapted algorithms and software for data analysis; M.L., T.L., Y.C., M.G. and F.L. coordinated the acquisition, distribution and quality evaluation of samples; Y.Z. and H.Y. wrote the manuscript.

## Funding

This work was funded by grants from the National Key R&D Program of China (2022YFF0608401) and the National Natural Science Foundation of China (31901038).

## Notes

### Competing Interest Statement

The authors have declared no competing interest.

