## Supplementary Information for "Proteomic Landscape of Human Spermatozoa: Optimized Extraction Method and Application"

**Running title: Human Spermatozoa Protein Profiling After** **Antibiotic Therapy**

***Corresponding Authors:**

Yong Zhang, PhD, Associate Researcher, Institutes for Systems Genetics, West China Hospital, Sichuan University, Chengdu, China;

Fuping Li, PhD, Associate Professor, Human Sperm Bank, West China Second University Hospital, Sichuan University, Chengdu, China

**Address:** No. 1, Keyuan 4th Road, Gaopeng Avenue, Hi-tech Zone, Chengdu 610041, China. Phone: +86-28-85164031; Fax: +86-28-85164031;

**Table of Contents**

**Figure. S1.** Heatmap analysis of different proteins between UA_Ultrasonication and SDS_Ultrasonication methods

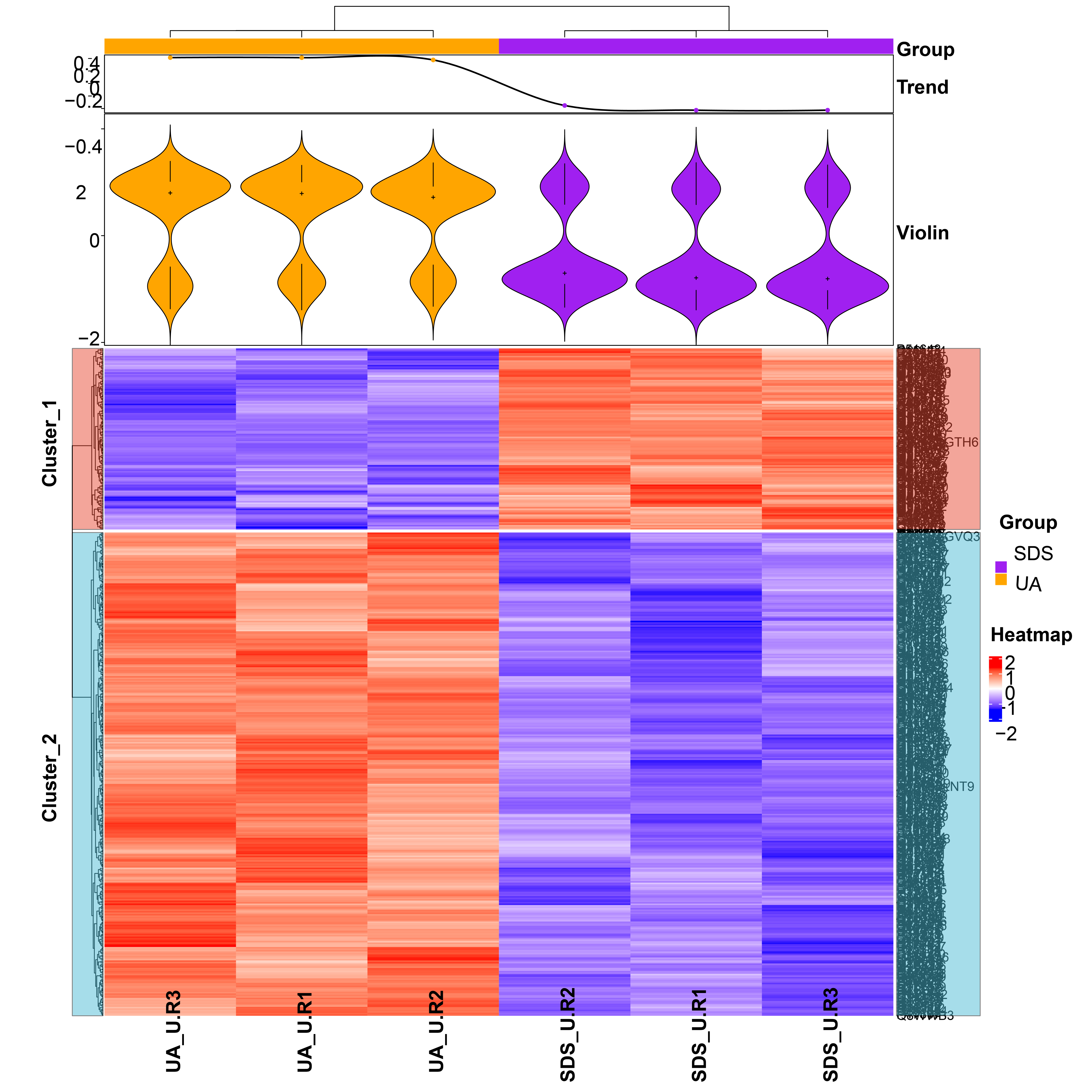

**Figure S2.** GO analysis of different proteins between UA_Ultrasonication and SDS_Ultrasonication methods

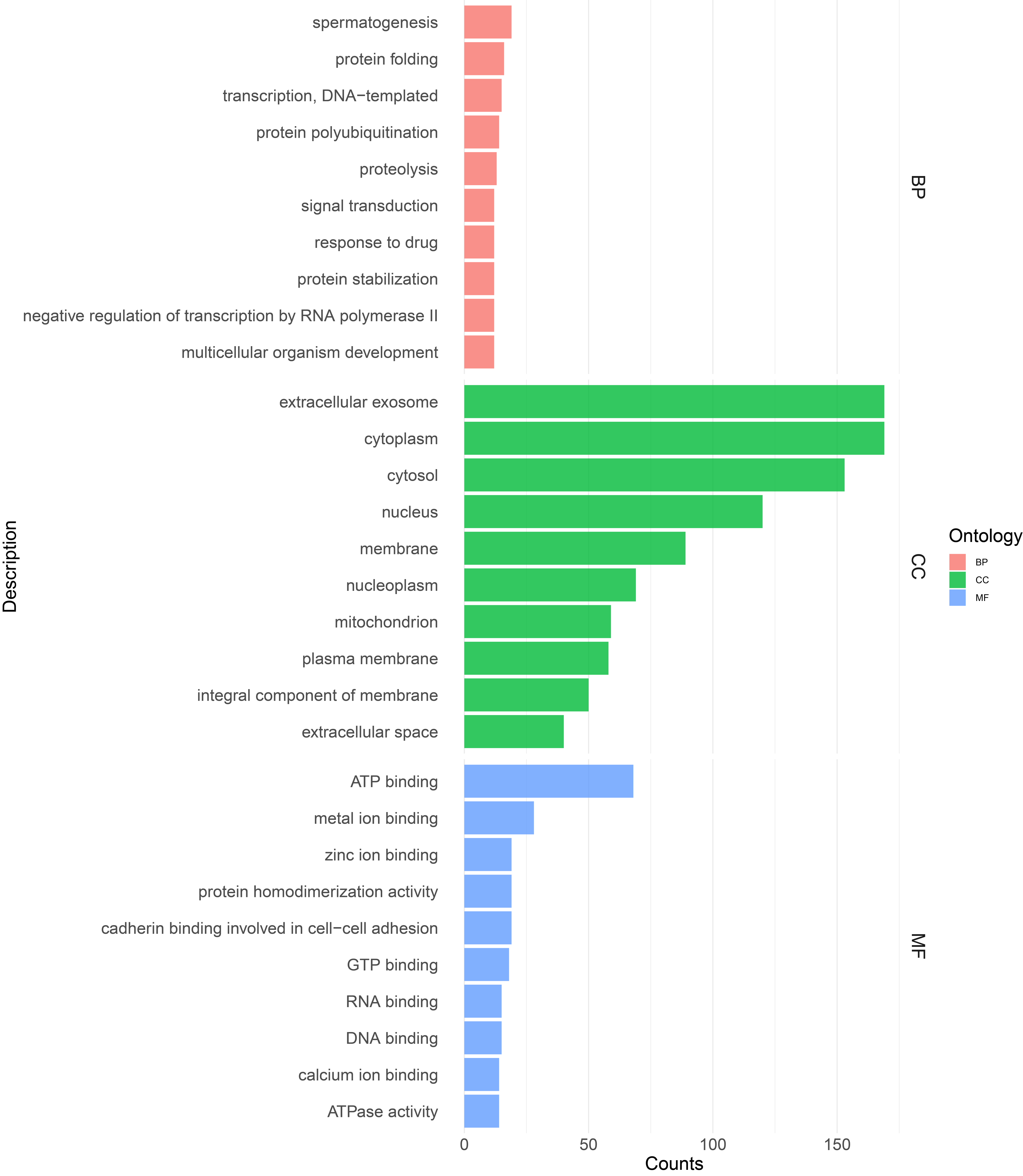
